# *Fendrr* synergizes with Wnt signalling to regulate fibrosis related genes during lung development via its RNA:dsDNA Triplex Element

**DOI:** 10.1101/2021.11.02.466973

**Authors:** Tamer Ali, Sandra Rogala, Nina M. Krause, Jasleen Kaur Bains, Maria-Theodora Melissari, Sandra Währisch, Harald Schwalbe, Bernhard G Herrmann, Phillip Grote

**Affiliations:** Institute of Cardiovascular Regeneration, Centre for Molecular Medicine, Goethe University, Theodor-Stern-Kai 7, 60590 Frankfurt am Main, Hesse, Germany; Faculty of Science, Benha University, Benha 13518, Egypt; Department of Developmental Genetics, Max Planck Institute for Molecular Genetics, Ihnestr. 63-73, 14195 Berlin, Germany; Center for Biomolecular Magnetic Resonance (BMRZ), Institute for Organic Chemistry and Chemical Biology, Goethe University, Max-von-Laue-Str. 7, 60438, Frankfurt am Main, Hesse, Germany; Georg-Speyer-Haus, Paul-Ehrlich-Str. 42-44, 60596 Frankfurt am Main, Hesse, Germany

## Abstract

Long non-coding RNAs are a very versatile class of molecules that can have important roles in regulating a cells function, including regulating other genes on the transcriptional level. One of these mechanisms is that RNA can directly interact with DNA thereby recruiting additional components such as proteins to these sites via a RNA:dsDNA triplex formation. We genetically deleted the triplex forming sequence (*FendrrBox*) from the lncRNA *Fendrr* in mice and find that this *FendrrBox* is partially required for *Fendrr* function *in vivo*. We find that the loss of the triplex forming site in developing lungs causes a dysregulation of gene programs, associated with lung fibrosis. A set of these genes contain a triplex site directly at their promoter and are expressed in fibroblasts. We confirm the formation of RNA:dsDNA formation with target promoters. We find that *Fendrr* with the Wnt signalling pathway regulates these genes, implicating that *Fendrr* synergizes with Wnt signalling in lung fibrosis.

## INTRODUCTION

The number of loci in mammalian genomes that produce RNA that do not code for proteins is higher than the number of loci that produce protein coding RNAs (1,2). These non-protein coding RNAs are commonly referred to long non-coding RNAs (lncRNAs) if their transcript length exceeds 200 nucleotides. Many of these lncRNA loci are not conserved across species. However, some loci are conserved on the syntenic level and some even on the transcript level. One of the syntenic conserved lncRNAs is the *Fendrr* gene, divergently expressed from the essential transcription factor coding gene *Foxf1*. Both genes have been implicated in various developmental processes (3–5) and particularly in heart and lung development (6–9).

The *Fendrr* lncRNA was shown to be involved in several pathogeneses with fibrotic phenotypes. In a transverse aortic constriction (TAC) mouse model, *Fendrr* was upregulated in heart tissue. Loss of *Fendrr* RNA via an siRNA approach alleviated fibrosis induced by TAC, demonstrating a pro-fibrotic function for *Fendrr* in the heart (10). In contrast, in humans with Idiopathic Pulmonary Fibrosis (IPF) and in mice with bleomycin-induced pulmonary fibrosis, the *Fendrr/FENDRR* RNA was downregulated (11). In addition, depletion of *FENDRR* increases cellular senescence of human lung fibroblast. While overexpression of human *FENDRR* in mice reduced bleomycin-induced lung fibrosis, revealing an anti-apoptotic function of *FENDRR* in lungs and a conservation of the mouse Fendrr and the human *FENDRR* in this process. *Fendrr/FENDRR* seems to have opposing functions on fibrosis in heart and in lung tissue, indicating that secondary cues such as active signalling pathways might be required.

In the lung, *FENDRR* is a potential target for intervention to counteract fibrosis and the analysis of its function in this process and how target genes are regulated is of interest to develop RNA-based therapies (12). LncRNAs can exert their function on gene regulation via many different mechanisms (13). One mechanism is that the RNA is tethered to genomic DNA either by base-pairing or by RNA:dsDNA triplex formation involving Hoogsteen base pairing (14). Here, we deleted such a Triplex formation site in the *Fendrr* lncRNA *in vivo*. We identified genes that are regulated by *Fendrr* in the developing mouse lung and require the triplex forming RNA element, which we termed the *FendrrBox*.The gene network that is regulated by *Fendrr* and require the *FendrrBox* element is associated with extracellular matrix deployment and with lung fibrosis. We verified that regulation of these genes is depending on the presence of full length *Fendrr* and active Wnt signalling, establishing that *Fendrr*, and, in particular, its *FendrrBox* element, is involved in Wnt dependent lung fibrosis.

## MATERIAL AND METHODS

### Culturing of mouse ES cells

The mESC were either cultured in feeder free 2i media or on feeder cells (mitomycin inactivated SWISS embryonic fibroblasts) containing LIF1 (1000 U/ml). 2i media: 1:1 Neurobasal (Gibco #21103049) :F12/DMEM (Gibco #12634-010), 2 mM L-glutamine (Gibco), 1x Penicillin/ Streptomycin (100x penicillin (5000 U/ml,) / streptomycin (5000ug/ml), Sigma #P4458-100ML, 2 mM glutamine (100x GlutaMAX™ Supplement, Gibco #35050-038), 1x non-essential amino acids (100x MEM NEAA, Gibco #11140-035), 1x Sodium pyruvate (100x, Gibco, #11360-039), 0.5x B-27 supplement, serum-free (Gibco # 17504-044), 0.5x N-2 supplement (Gibco # 17502-048), Glycogen synthase kinase 3 Inhibitor (GSK-Inhibitor, Sigma, # SML1046-25MG), MAP-Kinase Inhibitor (MEK-Inhibitor Sigma, #PZ0162), 1000 U/ml Murine_Leukemia_Inhibitory_Factor ESGRO (10^7^ LIF, Chemicon #ESG1107), ES-Serum media: Knockout Dulbecco’s Modified Eagle’s Medium (DMEM Gibco#10829-018), ES cell tested fetal calf serum (FCS), 2 mM glutamine, 1x Penicillin/ Streptomycin, 1x non-essential amino acids, 110 nM ß-Mercaptoethanol, 1x nucleoside (100x Chemicon #ES-008D), 1000 U/ml LIF1.

The cells were split with TrypLE Express (1x, Gibco #12605-010) and the reaction was stopped with the same amount of Phosphate-Buffered Saline (PBS Gibco #100100239) followed by centrifugation at 1000 rpm for 5min. The cells were frozen in the appropriate media containing 10% Dimethyl sulfoxide (DMSO, Sigma Aldrich #D5879). To minimize any effect of the 2i (Choi et al., 2017) on the developmental potential mESC were only kept in 2i for the antibiotic selection after transient transfection with CRISPR/Cas9 or mini gene integration and DNA generation for genotyping. At all other times cells were maintained on ES-Serum media on feeder cells.

### Generation of transgenic or CRISPR/Cas9 edited mESC

Guide RNAs were designed, using the crispr.mit.edu website with the nickase option. The following, top-scoring guide RNAs were selected and cloned into pX330 (Addgene, # 42230) plasmid to allow for transient puromycin selection after transfection. The sgRNAs used for the deletion of the FendrrBox are upstream(L): TCAGGCAACACTCACTGGAC, downstream(R): GGGAAGACATGGGGGAGTAA. Wildtype F1G4 cells were transiently transfected with 2μg/mL puromycin (Gibco, #10130127) for 2 days and 1μg/mL puromycin for 1 day. Single mESC clones were picked 7-8 days after transfection and plated onto 96-well synthemax (Sigma, #CLS3535) coated plates and screened for genomic DNA deletion by PCR using primers outside of the deletion region.

### Genotyping of *Fendrr^3xpA/3xpA^* and *Fendrr^em7Phg/em7Phg^* tissues

The REDExtract-N-Amp™ Tissue PCR Kit (Merck, XNAT) was used for genotyping for all tissue explants. Genotyping of *FendrrNull* (*Fendrr^3xpA/3xpA^*) embryos with the three primers: Fendrr3xpA_F1: GCGCTCCCCACTCACGTTCC, Fendrr3xpA_Ra1: AGGTTCCTTCACAAAGATCCCAAGC, genoNCrna_Ra4: AAGATGGGGAACCGAGAATCCAAAG that will generate a 696bp band in wild type and a 371bp band when the 3xpA allele is present. Genotyping of *FendrrBox* (*Fendrr^em7Phg/em7Phg^*) embryo tissues with: FendrrBox_F2: ATGCTTCCAAGGAAGGACGG, FendrrBox_R2: CTTGACGCCAAGCTCCTGTA that generate a 602bp product in wild type and a 503bp product when the *FendrrBox* is missing.

### Lung preparation and RNA isolation

Staged embryo lungs were dissected from uteri into PBS and kept on ice in M2 media (Merck, M7167-50ML) until further processing. For direct RNA isolation the lung tissue was transferred into Precellys beads CK14 tubes (VWR, 10144-554) containing 1ml 900 μl Qiazol (Qiagen, #79306) and directly processed with a Bertin Minilys personal homogenizer. To remove the DNA 100 μl gDNA Eliminator solution was added and 180 μl Chloroform (AppliChem, #A3633) to separate the phases. The extraction mixture was centrifuge at full speed, 4°C for 15min. The aqueous phase was mixed with the same amount of 70 % Ethanol and transferred to a micro or mini columns depending of the amount of tissue and cells. The RNA was subsequently purified with the Qiagen RNAeasy Plus Min Kit (Qiagen, #74136) according to the manufacturers manual. Remaining tissue from the same embryos was used for genotyping to select homozygous mutants.

### Lung *ex vivo* culture

The lung culture was adopted from a previous published protocol (Hogan et al., 1994). Lungs were dissected from the E12.5 staged embryos in ice-cold PBS containing 0.5% FCS. Lungs were then placed in holding medium: Leibovitz’s L-15 Medium (ThermoFisher Scientific, 11415064) containing 1x Corning™ MITO+ Serum Extender (Fisher scientific, 10787521) and 1x Pen/Strep. Explant media (Advanced DMEM/F12 (ThermoFisher Scientific, 12634010), 5x Corning™ MITO+ Serum Extender, 1x Pen/Strep, 10% FCS) was placed into a 6-well tissue culture dish (0.8-1.0ml) and the 6-well plate fitted with Falcon™ Cell Culture Inserts with 8 um pore size (Fisher Scientific, 08-771-20). The lungs were transferred from the holding medium onto the membrane with a sterile razor blade and 5-10 ul of holding media to keep the lungs wet. Cells were cultured at 37C with an atmospheric CO_2_ of 7.5%. After the indicated time the lungs were removed from the membrane and RNA isolated as described above.

### Generation of mouse embryos from mESCs

All animal procedures were conducted as approved by local authorities (LAGeSo Berlin) under the license number G0368/08. Embryos were generated by tetraploid morula aggregation of embryonic stem cells as described in (George et al., 2007). SWISS mice were used for either wild-type donor (to generate tetraploid morula) or transgenic recipient host (as foster mothers for transgenic mutant embryos). All transgenic embryos and mESC lines were initially on a hybrid F1G4 (C57Bl6/129S6) background and backcrossed seven times to C57Bl6J for the preparations of embryonic lungs. The study is reported in accordance with ARRIVE guidelines (https://arriveguidelines.org/).

### Real-time quantitative PCR analysis

Quantitative PCR (qPCR) analysis was carried out on a StepOnePlus™ Real-Time PCR System (Life Technologies) using Fast SYBR™ Green Master Mix (ThermoFisher Scientific #4385612). RNA levels were normalized to housekeeping gene. Quantification was calculated using the ΔΔCt method (Muller et al., 2002). *Rpl10* served as housekeeping control gene for qPCR. The primer concentration for single a single reaction was 250nM. Error bars indicate the standard error from biological replicates, each consisting of technical triplicates. The Oligonucleotides for the qPCRs are as follows: Emp2_qPCR_fw: GCTTCTCTGCTGACCTCTGG, Emp2_qPCR_rv: CGAACCTCTCTCCCTGCTTG, Serpinb6b_qPCR_fw: ATAAGCGTCTCCTCAGCCCT, Serpinb6b_qPCR_rv: CTTTTCCCCGAAGAGCCTGT, Trim16_qPCR_fw: CCACACCAGGAGAACAGCAA, Trim16_qPCR_rv: AGGTCCAACTGCATACACCG, Fn1_qPCR_fw: GAGTAGACCCCAGGCACCTA, Fn1_qPCR_rv: GTGTGCTCTCCTGGTTCTCC, Akr1c14_qPCR_fw: TGGTCACTTCATCCCTGCAC, Akr1c14_qPCR_rv: GCCTGGCCTACTTCCTCTTC, Ager_qPCR_fw: TGGTCAGAACATCACAGCCC, Ager_qPCR_rv: CATTGGGGAGGATTCGAGCC, Fendrr_qPCR_fw: CTGCCCGTGTGGTTATAATG, Fendrr_qPCR_rv: TGACTCTCAAGTGGGTGCTG, Foxf1_qPCR_fw: CAAAACAGTCACAACGGGCC, Foxf1_qPCR_rv: GCCTCACCTCACATCACACA, Rpl10_qPCR_fw: GCTCCACCCTTTCCATGTCA, Rpl10_qPCR_rv: TGCAACTTGGTTCGGATGGA.

### Sequencing and analysis of RNA-seq

RNA was treated to deplete rRNA using Ribo-Minus technology. Libraries were prepared from purified RNA using ScriptSeq™ v2 and were sequenced on an Illumina HiSeq platform. We obtained 60 million paired-end reads of 50 bp length. Read mapping was done with STAR aligner using default settings with the option --outSAMtype BAM SortedByCoordinate (Dobin et al., 2013) with default settings. For known transcript models we used GRCm38.100 Ensembl annotations downloaded from Ensembl repository (Zerbino et al., 2018). Counting reads over gene model was carried out using GenomicFeatures Bioconductor package (Lawrence et al., 2013).The aligned reads were analyzed with custom R scripts in order to obtain gene expression measures. For normalization of read counts and identification of differentially expressed genes we used DESeq2 with Padj < 0.01 cutoff (Love et al., 2014). GO term and KEGG pathways were analysed using g:Profiler (Raudvere et al., 2019). The data are deposited to GEO and can be downloaded under the accession number GSE186703.

### Triplex prediction

To calculate *Fendrr* triplex targets, DE genes from *FendrrNull* and *FendrrBox* RNA-Seq output were intersected and RNA-DNA triplex forming potential of the shared genes were calculated with Triplex Domain Finder (TDF) algorithm (Kuo et al., 2019). The command was executed with promotertest option and –organism = mm10. The rest of the options were set to the default settings.

### Culturing of NIH3T3 cells

NIH3T3 cells were cultured in DMEM (Gibco #11960-044) containing 10% Bovine Serum (Fisher Scientific #11510526), 1% GlutaMAX™ (Gibco #35050-038) and 1% Penicillin-Streptomycin (Sigma Aldrich #P4458). For the experiment, the cells were detached using Trypsin-EDTA (Gibco #25300-054). The reaction was stopped by adding double the amount of fresh media followed by centrifugation at 1000 rpm for 4 min. The pellet was resuspended in fresh medium and counted using a Chemometec NucleoCounter NC-200 Automated Cell Counter (Wotol #2194080-18). 0.15 ×10^6^ cells were seeded per well (Greiner Bio-One™ #657160).

### CRISPR-activation of *Fendrr* and treatment of NIH3T3 cells

Three guide RNAs targeting the *Fendrr* promoter were designed using the crispor.tefor.net website (Concordet and Haeussler, 2018). *Fendrr_sg1:* GGCCTCCGACGCTGCGCGCC, *Fendrr_sg2:*TCAACGTAAACACGTTCCGG, *Fendrr_sg3:* AGTTGGCCTGATGCCCCTAT. A non-specific guide RNA *ctrl_sg:* GGGTCTTCGAGAAGACCT served as control. The guide RNAs were cloned into the sgRNA(MS2) plasmid (addgene #61424). The CRISPR SAM plasmid (pRP[Exp]-Puro-CAG-dCAS9-VP64:T2A:MS2-p65-HSF1) was a gift from Mohamed Nemir from the Experimental Cardiology Unit Department of Medicine University of Lausanne Medical School.

For transfection, Lipofectamine 3000 (Invitrogen #L3000001) was used following the manufacturer’s guidelines. Briefly, 1μg total plasmid DNA (1:3 SAM to gRNA ratio) was diluted in Opti-MEM (Gibco #31985062) and mixed with p3000 reagent. Lipofectamine reagent was diluted in Opti-MEM and subsequently added to the DNA mixture. During the incubation the cells were washed with DPBS (Gibco #14190250) and provided with fresh Opti-MEM. Transfection mix was added to the cells and incubated for 4h at 37 °C. After the incubation, the media was changed with full media containing FGF (10 ng/ml bFGF, Sigma Aldrich #F0291; 25 ng/ml rhFGF, R&D Systems #345-FG), BMP-4 (40 ng/ml, R&D Systems #5020-BP-010) or CHIR99021 (3 μM, Stemcell #72052). The treatment was replenished by changing media after 24h, cells were harvested for RNA isolation after 48h.

### CD spectroscopy and melting curve analysis

Circular dichroism spectra were acquired on a Jasco J-810 spectropolarimeter. The measurements were recorded from 210 to 320 nm at 25 °C using 1 cm path length quartz cuvette. CD spectra were recorded on 8 μM samples of each DNA duplex and RNA:dsDNA triplex in 20 mM LiOAc, 10 mM MgCl_2_, pH 5.5. For the RNA:dsDNA triplex an excess of 5 eq RNA was used. By hybridization DNA duplex and RNA:dsDNA triplex were formed. Therefore, the complementary DNA strands were incubated at 95 °C for 5 min and afterwards cooled down to room temperature. For the triplex formation RNA was added to the DNA duplex and incubated at 60 °C for 1 h and then cooled down to room temperature (15). The sequences used are listed below. Spectra were acquired with 8 scans and the data was smoothed with Savitzky-Golay filters. Observed ellipticities recorded in millidegree (mdeg) were converted to molar ellipticity [θ] = deg x cm^2^ x dmol^-1^. Melting curves were acquired at constant wavelength using a temperature rate of 1 °C/min in a range from 5 °C to 95 °C. All melting temperature data was converted to normalized ellipticity and evaluated by the following equation using SigmaPlot 12.5:

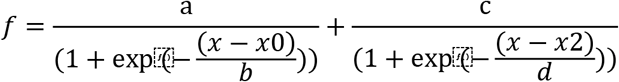

The RNA and DNA oligos used here were: Fendrr_Short: UCUUCUCUCUCCUCUCUUCUCCCUCCCCUC (30 nt), Fendrr_Long: UCUUCUCUCUCCUCUCUUCUCCCUCCCCUCCAUCCUCUUCCUUCUCCUCCUCCUCUU (57 nt), Emp2 (GA-rich): AGGAGAGAGAGGAGAGAGGGGAGAGAGGGG (30 nt), Emp2 (CT-rich): CCCCTCTCTCCCCTCTCTCCTCTCTCTCCT (30 nt), Foxf1 (GA-rich): CCGAGCCGGGAGGAGGAGGAGGAGCAGGAGGGGAGGGAGGGGAGGGGGCT (50 nt): Foxf1 (CT-rich): AGCCCCCTCCCCTCCCTCCCCTCCTGCTCCTCCTCCTCCTCCCGGCTCGG (50 nt)

## RESULTS

### The *FendrrBox* region is partially required for *Fendrr* RNA function

We established previously that the long non-coding RNA *Fendrr* is an essential lncRNA transcript in early heart development in the murine embryo (3). In addition, the *Fendrr* locus was shown to play a role in lung development (7). Expression profiling of pathological human lungs revealed that FENDRR is dysregulated in disease settings (16). In the second to last exon of the murine *Fendrr* lncRNA transcript resides a UC-rich low complexity region of 38bp, which can bind to target loci and thereby tether the *Fendrr* lncRNA to the genome of target genes(17). To address if this region is required for *Fendrr* function, we deleted this *FendrrBox* (*Fendrr^em7Phg/em7Phg^*) in mouse embryonic stem cells (mESCs) (Figure 1A). We generated embryos from these mESCs and compared them to the *Fendrr null* phenotype (Figure 1B). The *FendrrNull* (*Fendrr^3xpA/3xpA^*) embryos exhibit increased lethality starting at the embryonic stage E12.5 and all embryos were dead prior to birth and in the process of resorption (3). In contrast, the *FendrrBox* mutants, which survived longer, displayed an onset of lethality later during development (E16.5) and some embryos survived until short before birth. The surviving embryos of the *FendrrBox* mutants were born and displayed an increased postnatal lethality (Figure 1C). This demonstrates that the *FendrrBox* element in the *Fendrr* RNA is most likely partially required for *Fendrr* function in embryo development and for postnatal survival.

**Figure 1.**
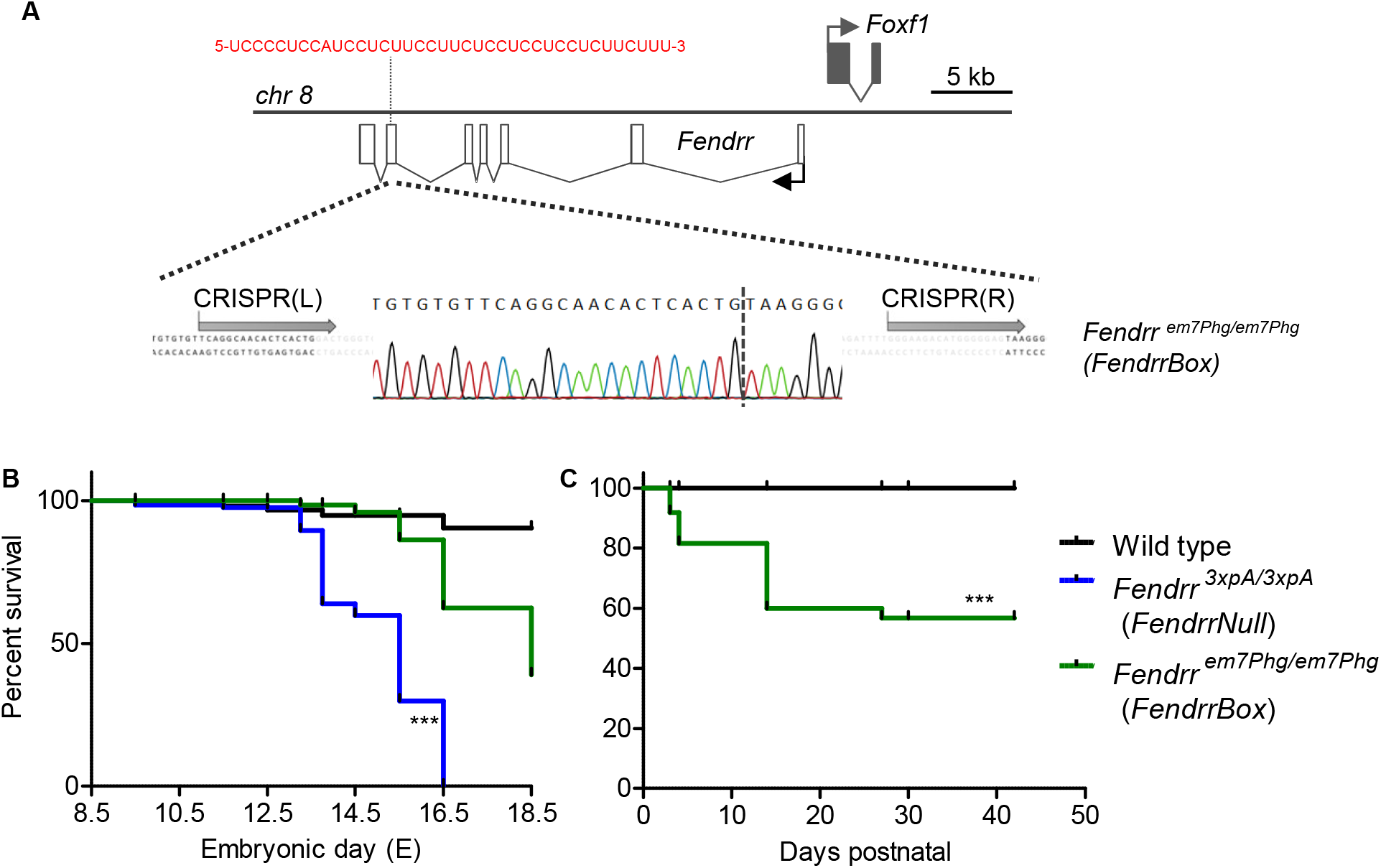
(**A**) Schematic of the *Fendrr* locus and the localization of the DNA interacting region (*FendrrBox*) in exon six. The localization of the gRNA binding sites (grey arrows) are indicated and the resulting deletion of 99bp, including the *FendrrBox*, in the genome that generates the *Fendrr^em7PhG/em7PhG^* allele (*FendrrBox*). (**B**) Embryos and (**C**) life animals were generated by tetraploid aggregation and the surviving animals counted. *** p>0.0001 by logrank (Mantel-Cox) test

### Gene expression in *FendrrNull* and *FendrrBox* mutant developing lungs

Given the involvement of *Fendrr* in lung development (7) and the involvement of mutations in human *FENDRR* in lung disease (9), we wanted to determine the genes affected by a loss of *Fendrr* or the *FendrrBox* in developing lungs to identify *Fendrr* target genes. However, when we collected the lungs from surviving embryos of the E14.5 stage we did not identify any significant dysregulation of genes, neither in the *FendrrNull* nor in the *FendrrBox* mutant lungs (Figure S1). One explanation is that the incomplete genetic penetrance of *Fendrr* mutants results in a compositional bias. Surviving embryos do not display any differences in gene regulation and those which did, were lethal and the embryos died already. To circumvent this issue, we collected embryonic lungs from E12.5 stage embryos, before the timepoint that any lethality occurs and cultivated the lung explants *ex vivo* under defined conditions. After 5 days of cultivation some lungs from all phenotypes detached from the supporting membrane. Hence, we choose to analyze 4 day cultivated lungs (corresponding then to E16.5) (Figure 2A). When we compared expression between wild type, *FendrrNull* and *FendrrBox* mutant E16.5 *ex vivo* lungs we found 119 genes dysregulated in *Fendrr null* and 183 genes in *FendrrBox* mutant lungs compared to wild type (Figure 2B). When we analysed the GO terms of downregulated genes in both *Fendrr* mutants we found mainly genes involved in lung and respiratory system development, as well as cell-cell contact organization and extracellular matrix organization (Figure 2C). Upregulated genes in both mutants were mostly associated with genome organization, replication, and genome regulation. Overall, 60 genes were commonly dysregulated in both mutants (Figure 2D). Strikingly, these shared dysregulated genes are mostly associated with lung fibrosis, a major condition of various lung disease, including idiopathic pulmonary fibrosis (IPF).

**Figure 2.**
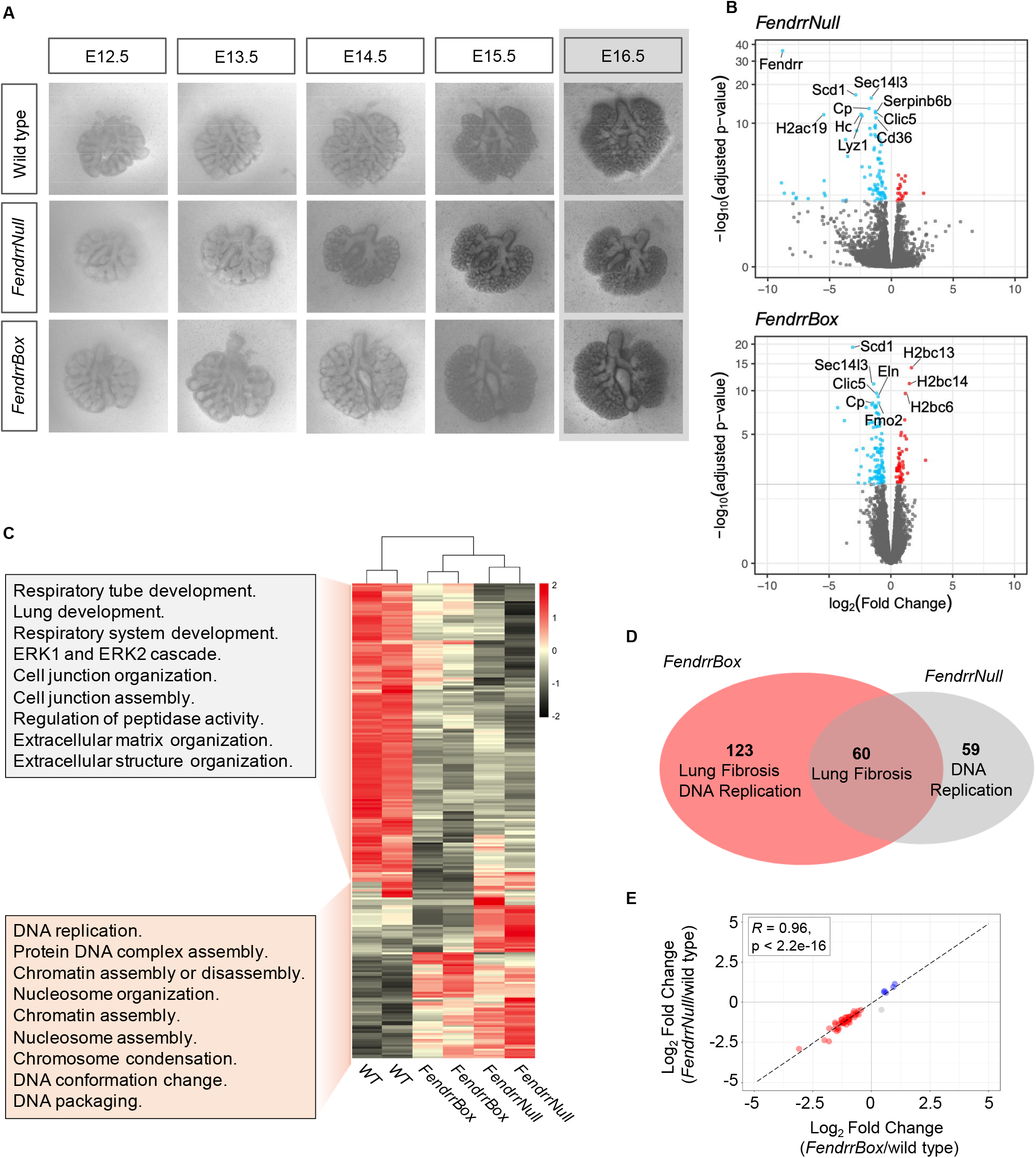
Expression profiling of *Fendrr* mutant lungs in *ex vivo* development. (**A**) Representative images from a time course of *ex vivo* developing lungs from the indicated genotype. The last time point representing E16.5 of embryonic development is the endpoint and lungs were used for expression profiling. (**B**) Volcano plot representation of deregulated genes in the two *Fendrr* mutants determined by RNA-seq of two biological replicates. (**C**) Heatmap of all 242 deregulated genes of both *Fendrr* mutants compared to wild type. The GO terms of the either up- or downregulated gene clusters are given in the box as determined by topGO bioconductor package. (**D**) Venn diagram of the individually deregulated genes and the overlap in the two different *Fendrr* mutants. Pathway analysis performed by wikiPathaways is given for each DE genes cluster. (**E**) Log_2_ fold change scatter plot of the 60 DE genes shared between *FendrrBox* and *FendrrNull*. The correlation coefficient between the log_2_ fold change is calculated using Pearson correlation coefficient test. Blue and red indicate that genes are upregulated and downregulated in both mutants, respectively, whereas gray indicates the that genes are changing in either mutant.

### RNA:dsDNA triplex target genes in fibrosis

It is conceivable that some of these dysregulated genes are primary targets of *Fendrr* and some represent secondary targets. To identify which of these dysregulated genes in *Fendrr* mutant lungs are likely to be direct targets of *Fendrr* via its triplex forming *FendrrBox*, we used the Triplex Domain Finder (TDF) algorithm (18) to identify triplex forming sites on *Fendrr* within the promoters of the dysregulated target genes. The single significant triplex forming site (or DBD = DNA Binding Domain) discovered by TDF is the *FendrrBox* (Figure 1A, 3A), confirming previous results. The TDF algorithm didn’t identify significant binding of *Fendrr* to target promoters in either *FendrrBox* exclusive nor the *FendrrNull* exclusive dysregulated genes. However, the TDF algorithm detects a significant *FendrrBox* binding site in promoters of 20 out of the 60 target genes from the overlapping gene set of *FendrrBox* and *FendrrNull* mutants (Figure 3A). We refer to these genes as direct *FendrrBox* target genes and most of these 20 genes are downregulated in loss of function *Fendrr* mutants (Figure 3B). When we analyzed more closely the GO terms associated with these shared genes, we find most terms to be associate with cell adhesion and extracellular matrix functions, a typical hallmark for fibrosis, where collagen and related components are deposited from cells (Figure S2).

**Figure 3.**
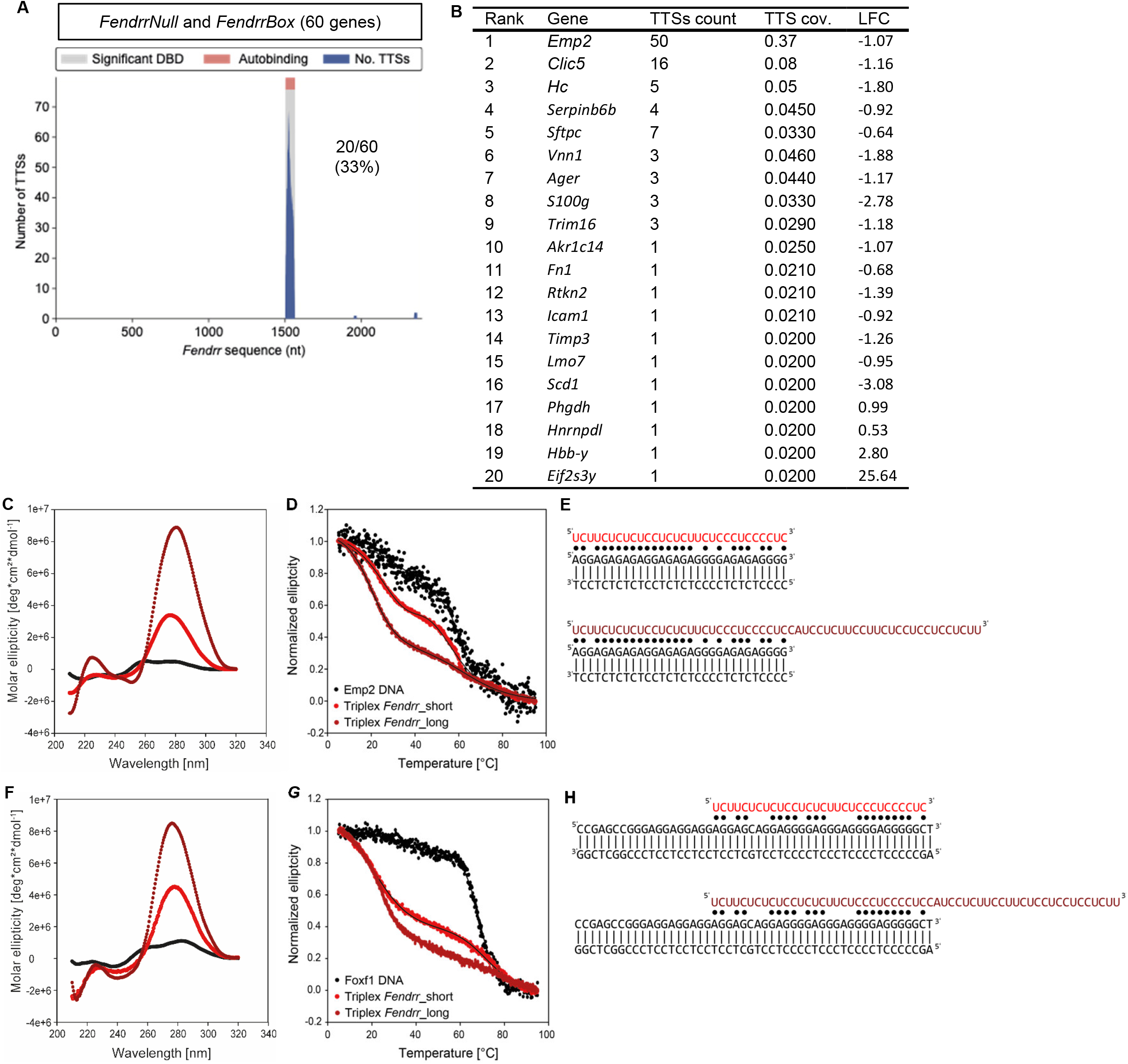
Potential direct target genes of *Fendrr* and RNA:dsDNA triplex formation capacity. (**A**) Triplexes prediction analysis of the 60 shared dysregulated genes identifies 20 genes with a potential *Fendrr* triplex interacting site at their promoter. DBD = DNA Binding Domain on RNA, TTS = triple target DNA site. (**B**) List of the 20 *Fendrr* target genes that depend on the *Fendrr* triplex and have a *Fendrr* binding site at their promoter. (**C**) Circular dichroism spectra of the *Emp2:Fendrr_short* triplex (light red), the Emp2 DNA duplex (black) and the *Fendrr_long:*Emp2 triplex (dark red) measured at 298 K. (**D**) Thermal melting of the *Fendrr_short:Emp2* triplex (light red), the Emp2 DNA duplex (black) *Fendrr_long:*Emp2 triplex (dark red). (**E**) Sequence of Emp2 DNA duplex (black) *Fendrr_short* RNA (light red) *Fendrr_long* RNA (dark red), Watson-Crick base pairing is indicated with | and the Hoogsteen base pairing is indicated with ●. (**F**) Circular dichroism spectra of the *Fendrr_short:*Foxf1 triplex (light red), the *Foxf1* DNA duplex (black) and the *Fendrr_long:Foxf1* triplex (dark red) measured at 298 K. (**G**) Thermal melting of the *Foxf1:Fendrr_short* triplex (light red), the *Foxf1* DNA duplex (black) and the *Fendrr_long:Foxf1* triplex (dark red). (**H**) Sequence of *Foxf1* DNA duplex (black) *Fendrr_short* RNA (light red) and of *Fendrr_long* RNA (dark red), Watson-Crick base pairing is indicated with | and the Hoogsteen base pairing is indicated with ●.

### Biophysical RNA:dsDNA triplex characterization

We used CD spectroscopy to confirm the predicted TDF triplex formation and used *Emp2* as target gene for *Fendrr*. The CD spectrum presented typical features for triplex formation including an increased peak at ~280 nm, a transition at ~260 nm, two negative peaks at ~240 nm and ~210 nm and a positive peak at ~220 nm (19,20) that differed from the *Emp2* DNA CD duplex spectrum (Figure 3C). Moreover, we performed thermal melting assays to obtain the temperatures T_m_(DNA duplex) = 47.5 ± 2.6 °C, T_m1_(RNA:dsDNA _long) = 20.1 ± 0.1 °C and a second broad melting transition T_m2_(RNA_long:dsDNA) around 62 °C and T_m1_(RNA_short:dsDNA) = 41.1 ± 0.1 °C and a second melting transition T_m2_(RNA_short:dsDNA) = 59.6 ± 0.3 °C (Figure 3D). A unique feature of the triplex that we were able to characterize is the biphasic melting transition that results from the dissociation of the weaker Hoogsteen base pairs at the first melting point, followed by the melting of the Watson-Crick base pairs at the higher melting transition (Figure 3E).

We also investigated a previously predicted target gene *Foxf1* and *Fendrr* RNA triplex forming potential by CD spectroscopy. The triplex spectrum showed similar features as described before for the *Fendrr* RNA:Emp2 dsDNA triplex. The CD spectrum showed an increased negative ellipticity at ~210 nm and ~240 nm and positive ellipticity at ~280 nm (Figure 3F). Further, thermal melting data indicates features of triplex formation. The Triplex with the *Fendrr*_short RNA showed two melting points one T_m1_(RNA_short:dsDNA) = 20.9 ± 0.2 °C and a second broad melting transition T_m2_(RNA_short:dsDNA) around 72 °C and with the *Fendrr*_long RNA T_m1_(RNA_long:dsDNA) = 22.9 ± 0.1 °C and a second broad melting transition T_m2_(RNA_long:dsDNA) around 75 °C. For the duplex melting temperature T_m_(DNA duplex) = 68.8 ± 0.1 °C (Figure 3G). The data indicate the triplex forming potential of *Fendrr* RNA with different TDF predicted DNA sequences (Figure 3E, H).

### Signalling dependent regulation by *Fendrr*

To functionally test for direct *Fendrr* targets, we wanted to analyze the expression of these 20 genes in NIH3T3 mouse fibroblasts. Only 6 out of these 20 are expressed in this cell line and *Fendrr* is only very lowly expressed. To activate endogenous *Fendrr* expression we tested several gRNAs to recruit the dCAS9-SAM transcriptional activator complex (21) to the promoter region of *Fendrr*. We identified three gRNAs (Figure 4A) that could exclusively activate endogenous *Fendrr* without significant activation of the *Foxf1* gene (Figure 4B). Such transfected fibroblasts have a 15-fold increase in *Fendrr* transcript. Upon over activation of endogenous *Fendrr*, none of the expressed *FendrrBox* target genes displayed an increase in expression (Figure 4C), as it would be expected as these genes are downregulated in *Fendrr* loss-of-function mutants (Figure 3B). We speculated that in addition to overexpression of *Fendrr*, an additional pathway needs to be activated. The BMP, FGF and Wnt pathway are known to play an important role in lung fibrosis (22,23). We therefore activated the BMP signalling pathway, FGF-signalling pathway and the Wnt signalling pathway in these fibroblasts (Figure 4D-F). We found that only when Wnt signalling was activated, overactivation of *Fendrr* could increase the expression of nearly all the expressed *FendrrBox* target genes (Figure 4F). This places the lncRNA *Fendrr* as a direct coactivator of Wnt-signalling in fibroblasts and most likely in lung fibrosis.

**Figure 4.**
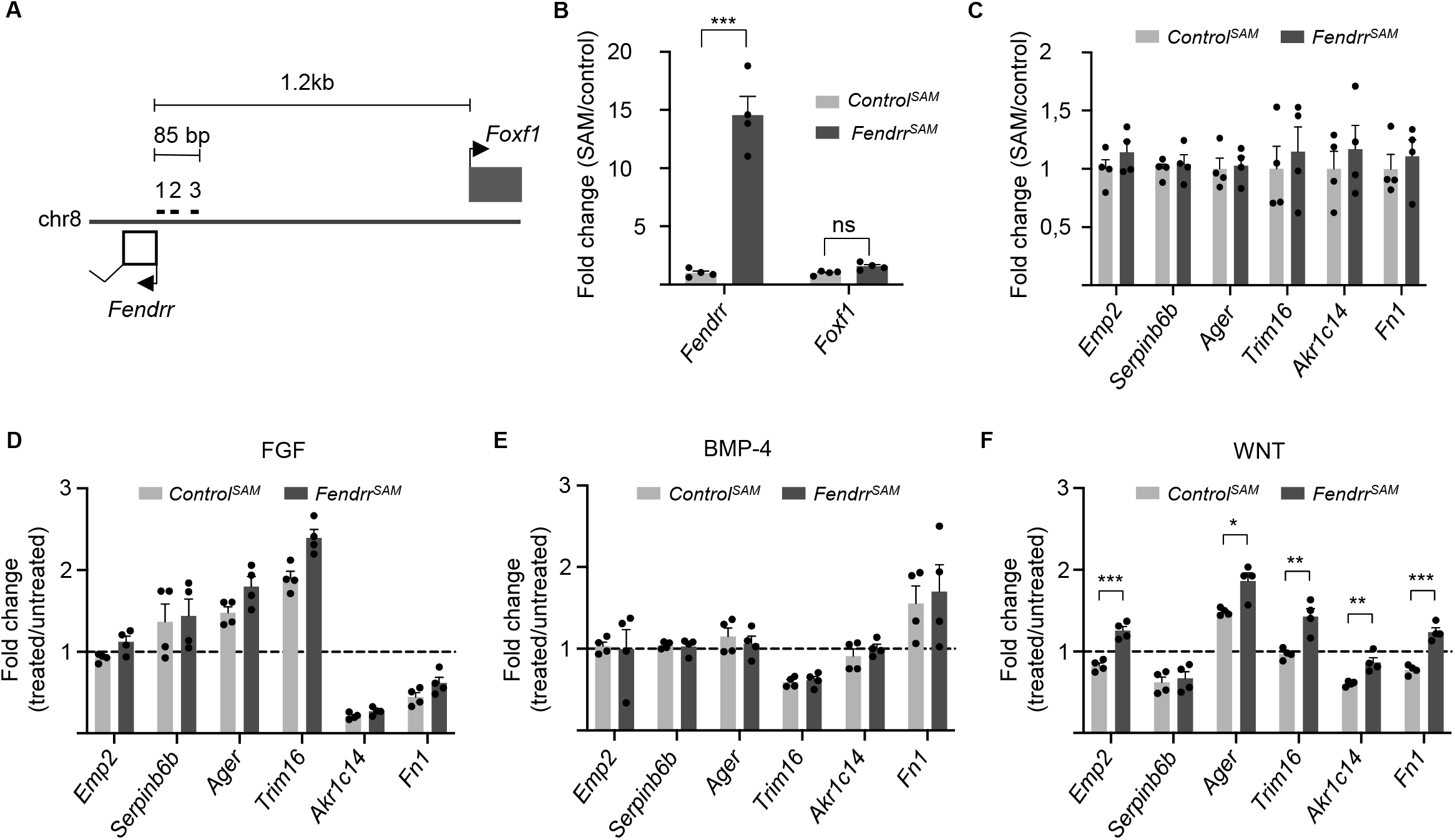
Wnt-dependent *Fendrr* target gene regulation. (**A**) Schematic of the *Foxf1* and *Fendrr* promoter region with the indication of the location of the 3 gRNAs used for specific *Fendrr* endogenous activation. (**B**) Increase of *Fendrr* expression in NIH3T3 cells upon CRISPRa with a pool of 3 gRNAs. (**C**) *Fendrr* Triplex containing *Fendrr* target genes expressed in NIH3T3 cells after 48h of *Fendrr* CRISPRa (*Fendrr^SAM^*). (**D**) Expression changes after 48hrs of co-stimulation with FGF. (**E**) Expression changes after 48hrs of co-stimulation with BMP-4. (**F**) Expression changes after 48hrs of co-stimulation with WNT. The dashed line represent the normalised expression value (set to 1) of the untreated cells transfected with control gRNA. (**D-F**) Statistics are given when significance by t-test analysis.

## DISCUSSION

We showed previously that *Fendrr* can bind to promoters of target genes in the lateral plate mesoderm of the developing mouse embryo (3,17). As *Fendrr* can also bind to histone modifying complexes, it is assumed that *Fendrr* directs these complexes to its target genes. However, that the *FendrrBox* might be the recruiting element was so far only supported by a biochemical approach that shows binding of the *FendrrBox* RNA element to two target promoters *in vitro* (3).

The involvement of *Fendrr* in lung formation was shown previously, albeit with a completely different approach to the removal of *Fendrr*. The replacement of the full length *Fendrr* locus by a *lacZ* coding sequence resulted in homozygous postnatal mice to stop breathing within 5h after birth (7). These mice also allowed for tracing *Fendrr* expression to the pulmonary mesenchyme, to which also vascular endothelial cells and fibroblasts belong. At the E14.5 stage *FendrrLacZ* mutant mice exhibit hypoplastic lungs. Our *ex vivo* analysis of lungs from our specific *Fendrr* mutants confirms the involvement of *Fendrr* in lung development. Here we show for the first time that the *FendrrBox* is at least partially required for *in vivo* functions of *Fendrr* and identified several, potential direct target genes of *Fendrr* in lung development. Moreover, the analysis of the dysregulated genes in the two different mouse mutant lungs indicates, that specifically *Fendrr* in the fibroblast might play an important role.

Studying embryonic development of the lung and its comparison to idiopathic lung fibrosis (IPF) in the adult lung has revealed that many of the same gene networks are in place to regulate both processes (24). A multitude of different signalling pathway are implicated in IPF (23). A prime example for an important pathway in IPF is the Wnt signalling pathway (25) and, in particular, increased Wnt signalling is associated with IPF and, hence, inhibition of Wnt signalling counteracts fibrosis (26). While the contribution of developmental signalling pathways to IPF is well understood, the contribution of lncRNAs in IPF is just beginning to be addressed (27). In humans, it was shown that in IPF patients *FENDRR* is increased in lung tissue (11). Intriguingly, in single cell RNA-seq approaches from human lung explants, *FENDRR* is highly expressed in vascular endothelial (VE) cells, but also significantly expressed in fibroblasts (28). Moreover, *FENDRR* expression increases in VE and in fibroblasts in IPF (28,29). It was shown recently, that *Fendrr* can regulate β-catenin levels in lung fibroblasts (30). Our data supports that Wnt signalling together with *Fendrr* is involved in target gene regulation and that *Fendrr* is a positive co-regulator of Wnt signalling in fibroblasts. This contrasts with the role of *Fendrr* in the precursor cells of the heart, the lateral plate mesoderm. Loss of *Fendrr* function results in the upregulation of *Fendrr* target genes, establishing that *Fendrr* is a suppressor of gene expression. The finding that *Fendrr* can act as either a suppressor or an activator of transcription, depending on the cell type, highlights the crosstalk between lncRNAs and signalling pathway, which broadens our understanding of the versatility of lncRNA in the cellular functions.

## DATA AVAILABILITY

The data are deposited to GEO and can be downloaded under the accession number GSE186703.

## SUPPLEMENTARY DATA

Supplementary Data are available at NAR online.

## ACKNOWLEDGEMENT

We thank Dijana Micic for excellent animal husbandry and Karol Macura for the generation of the transgenic mice. We want to thank Heiner Schrewe for help with ex vivo culture of embryonic lungs.

## FUNDING

This research was funded by the DFG (German Research Foundation) Excellence Cluster Cardio-Pulmonary System (Exc147-2) and a DFG research grant GR 4745/1-1 to P.G. T.A and S.R. are supported by the 403584255 – TRR 267 of the DFG.

## CONFLICT OF INTEREST

The authors declare no conflict of interest.

## Notes

### Competing Interest Statement

The authors have declared no competing interest.

### Summary of Updates

We added new authors to the manuscript and new experiments. The new experiments show that the predicted triplex site in Fendrr does form a triplex with target promoters in vitro. These data are added to Figure 3.

