## Supplementary material for "*Fendrr* synergizes with Wnt signalling to regulate fibrosis related genes during lung development via its RNA:dsDNA Triplex Element": S1

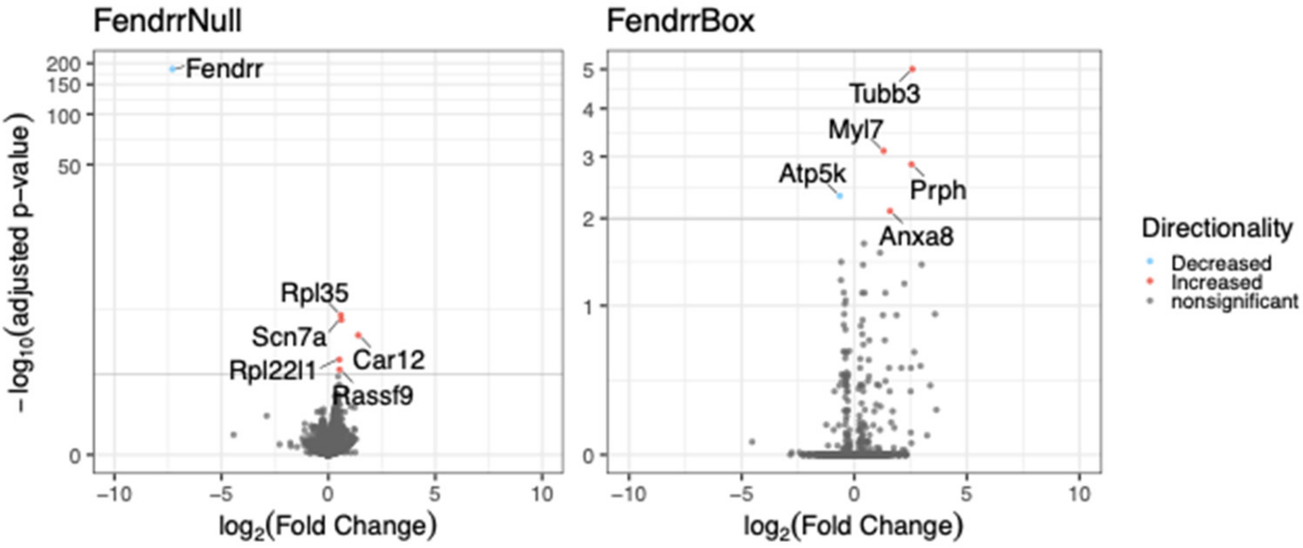


**Figure S1**. Expression profiling of *Fendrr* mutant E14.5 lungs in *in vivo* development.
Volcano plot representation of deregulated genes in the two *Fendrr* mutants determined by RNA-seq of three biological replicates.
