## Supplementary material for "*Fendrr* synergizes with Wnt signalling to regulate fibrosis related genes during lung development via its RNA:dsDNA Triplex Element": S2

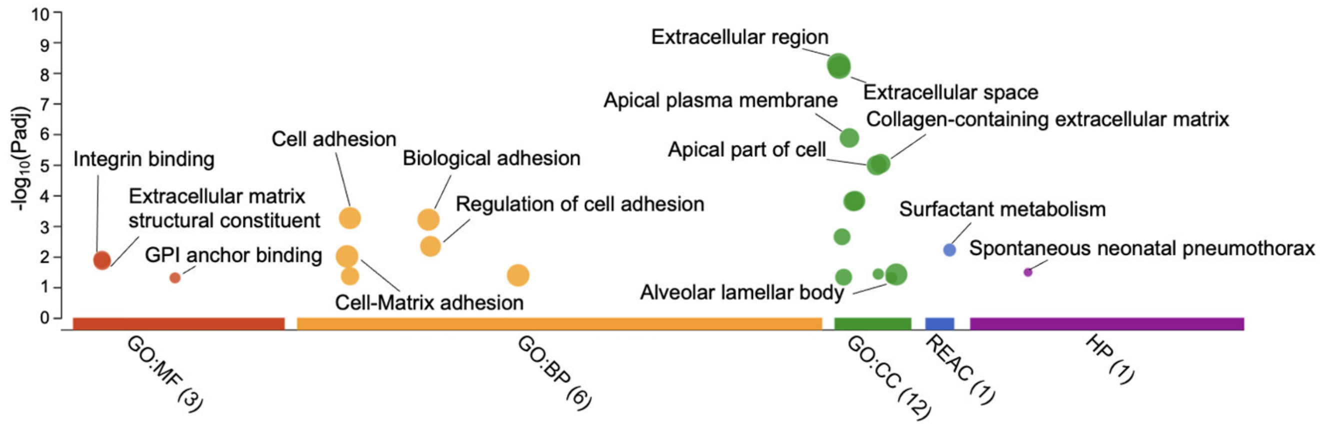


**Figure S2**. Functional Profiling of 20 direct *Fendrr* target genes

MF= molecular function, BP = biological process, CC = cellular component, Reactome (Reac), and Human pathways (HP) analysis. The size of each bubble represents the number of genes from the 20 genes that are involved in the enriched ontology.
